# Copy Number Variation in Gluthatione S-Transferase Variants using Multiplex Ligation-Dependent Probe Amplification in a Health Population in Goiânia – Go

**DOI:** 10.1101/384701

**Authors:** Lucas Carlos Gomes Pereira, Nádia Aparecida Bérgamo, Lucilene Arilho Bicudo, Jalsi Arruda Tacon, Bruno Faulin Gamba, Elisangela de Paula Silveira-Lacerda

## Abstract

Genetic polymorphisms in glutathione S-transferases (GSTs) genes might influence the detoxification activities of the enzymes predisposing individuals to a lot of diseases. Owing to the presence of these genetic variants, inter-individual and ethnic differences in GSTs detoxification capacity have been observed in various populations. Therefore, the present study was performed to determine GST variants in 100 healthy individuals from Goiânia – GO with a new methodology. GSTM1, GSTT1 and GSTP1 variants were analyzed by a MPLA (Multiplex Ligation by Probe Amplification) approach. The results obtained for the GSTM1 gene in exon 3 we found 43 deletions and 24 duplications, for exon 5 we found 45 deletions and 11 duplications, for the GSTT1 gene in exon 1 we found 14 deletions, and 18 duplications, for exon 5 we found 23 deletions and 28 duplications. For the GSTP1 gene in exon 3, we found only 1 deletion and for exon 4 we do not found any alteration. These findings in healthy population, give us such more information for the future epidemiological and clinical studies. Using to examine the effect of these combinations in drugs metabolism and cancer predisposition, further largest group would be needed, since their frequencies are quite low. To our of GSTs variations, this is the first study with this methodology indicating the frequencies of genetic variants of GST superfamily in a health population in a Goiânia population, and this study gives us new possibilities and data for new research using this same methodology.

## 1 Introduction

Individual inherited genetic differences related to polymorphism in detoxification enzymes could be an important factor not only in carcinogen metabolism but also in cancer susceptibility [1]. Functional genetic polymorphisms have been described for Glutathione-S-transferase (*GSTs*) genes, a superfamily of phase II metabolizing enzymes. GSTs catalyze the conjugation of reduced glutathione (GSH) to a wide variety of electrophilic compounds in order to make them more soluble enabling their elimination [2].

As a result of this detoxification activity, GSTs protect the cell from DNA damage, genomic instability and cancer development. In addition, as non-enzymatic proteins, GSTs can modulate signaling pathways that control cell proliferation, cell differentiation and apoptosis, among other processes [3, 4]. Differences in GSTs activity may modify the risk of cancer development and also may impact on the heterogeneous responses to toxic substances or specific therapies [2]. Moreover, GST mutations are known to contribute to in//ter-individual and ethnic variability in the susceptibility to environmental risk factors, cancer predisposition and drug responsiveness.

Several epidemiological studies evaluated the role of GST polymorphisms on CML susceptibility, but conflicting results have been achieved [5, 6]. CNVs are genomic that differ in copy number (CN) in compared genomes. CNVs include deletions, duplications, multiple duplications or more complex rearrangements. Common CNVs, also known as copy number polymorphisms (CNPs), account for approximately 10% of the human genome. Although CNVs are more frequent in intergenic regions, they overlap hundreds of protein-coding genes, regulatory sequences, and other functional genetic elements. Although the majority of CNVs are probably neutral, increasing numbers of CNVs are being associated with various human phenotypes, including diseases [7]. With the widespread use of substances (supplements, medicines, among others) and interaction with the environment and life style, there is a need to trace the metabolic genetic profile of people who practice physical activity to power end know and elucidate the that as changes found could affect the metabolism of these people. Therefore, the aim of the present study was investigate, for the first time, the importance of GST genetic mutations by CNV analysis in the health population in Goiânia - GO.

## 2 Methods

### 2.1 Subjects

This study was carried out in the Genetic Molecular and Citogenetic Laboratory of the Institute of Biological Sceince I at Federal University of Goias, Goiânia, Brazil. The study was performed acording approved by the ethics committee of Federal University of Goias (approval CEP: 895.552). In the current survey, 100 health people (35M/65F) of both genders and aged 18-90 (M: 30.2) years old were enrolled after receiving their informed consent.

### 2.2 Sample collection and DNA analyzes

Peripheral blood (5mL) was collected in EDTA vacutainer tubes from all participating individuals after obtaining their written consent. Genomic DNA extraction was performed from whole blood using Purelink DNA Kit (Invitrogen^®^).

### 2.3 Multiplex ligation-dependent probe amplification

Multiplex ligation-dependent probe amplification (MLPA) of *GSTM*, *GSTT* and *GSTP* were performed using the P128 CYP450 MLPA kit (Version C1; MRC-Holland^®^, Amsterdam, The Netherlands) according to the manufacturer’s instructions. In brief, all DNA samples were diluted to 200 ng with Tris-EDTA buffer and denatured in a thermalcycler for 5 min at 98°C. After cooling to 25°C, probemix and MLPA buffer were added to each sample, mixed and incubated for 1 min at 95°C followed by a 16 h hybridization at 60°C. The ligation reaction was performed at 54°C by adding 32 μl of Ligase-65 mix to each tube followed by heating for 5 min at 98°C. PCR buffer, water and MLPA ligation reactions were mixed in new tubes and maintained in a thermocycle at 60°C while polymerase mix was added to each tube. Exon-specific probes with universal tagged primers underwent PCR consisting of 35 amplification cycles (95°C for 30 s, 60°C for 30 s and 72°C for 60 s), followed by a 20 mini incubation at 72°C. Amplified products were separated by capillary gel electrophoresis ABI3500 ( Applied Biosystems^®^) and data were analyzed using *Coffalyser* v140721.1958 software (MRC-Holland^®^, The Netherlands).

## 3 Results

### 3.1 *GSTM*, *GSTT and GSTP* copy number analysis

To interrogate *GSTM1, GSTT1 and GSTP1* copy number, MLPA was performed on all 100 subjects DNA samples. The P128 Cytochrome P-450 MLPA kit (Version C1, MRC-Holland, The Netherlands) contained probes for *GSTM1, GSTT1 and GSTP1*. A representative MLPA analysis of 100 health people samples is shown in Figure 1.

**Figure 1.**
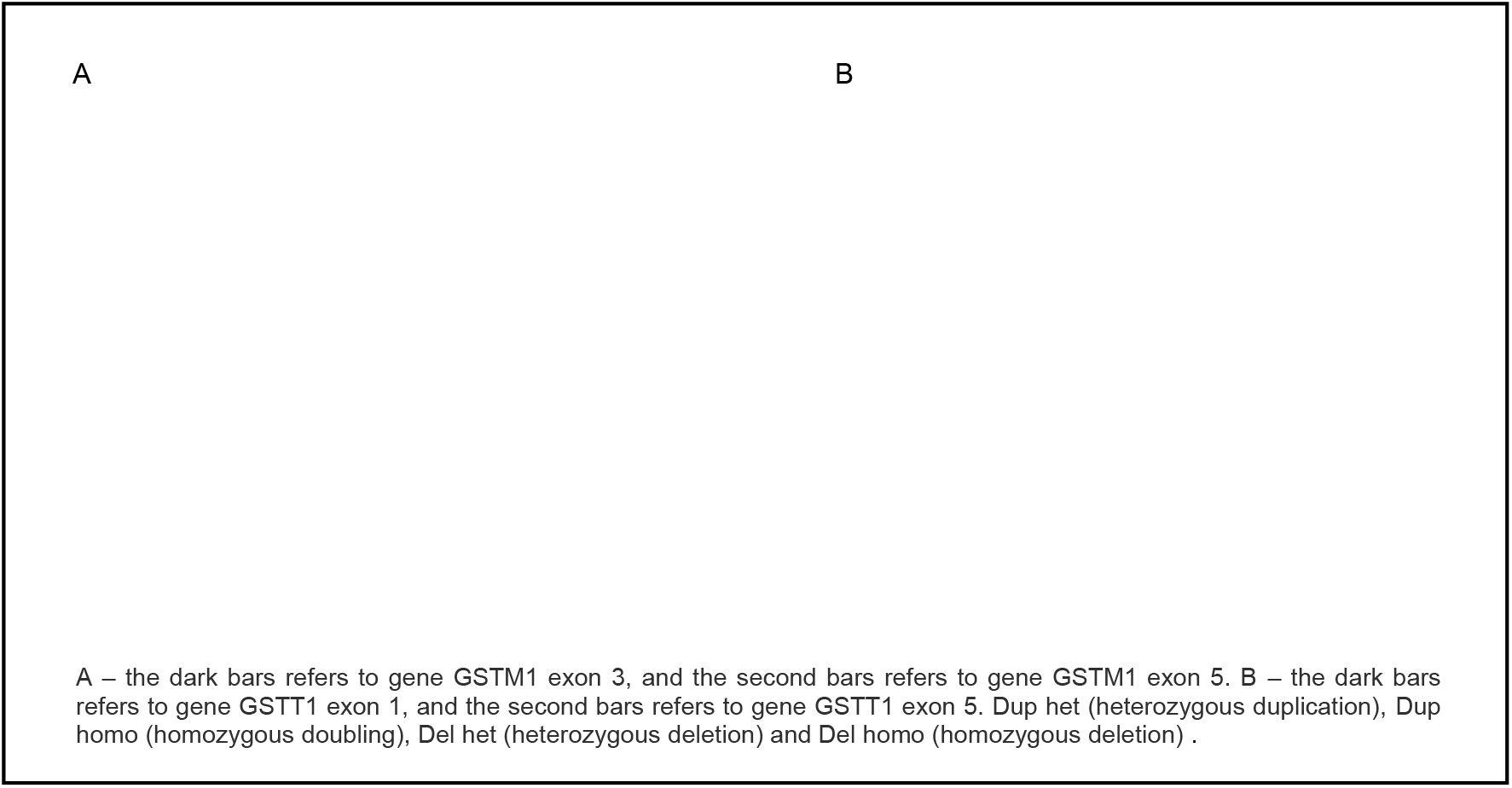
Representative *GSTM1* (exon 3 and exon 5) deletion and duplication and GSTT1 (exon 1 and exon 5) deletion and duplication analysis by MLPA.

For the *GSTM1* gene in exon 3 (Figure 1A) we found 43 deletions, all of which treat a deletion in homozygous (16 men and 26 women) and 24 duplications, being 14 homozygous (7 men and 7 women) and 10 heterozygous (2 men and 8 women). For exon 5 (Figure 1A) we found 45 deletions, being 43 homozygous (15 men and 28 women) and 2 heterozygous (1 man and 1 woman), and 11 duplications, being 10 homozygous (5 men and 5 women) and 1 heterozygous (1 woman).

For the *GSTT1* gene in exon 1 (Figure 1B) we found 14 deletions, being 13 homozygous (5 men and 8 women) and 1 heterozygous (1 woman), and 18 duplications, being 14 homozygous (8 men and 6 women) and 4 heterozygous (4 women). For exon 5 (Figure 1B) we found 23 deletions, being 20 homozygous (6 men and 14 women) and 3 heterozygous (3 women), and 28 duplications, being 21 homozygous (8 men and 13 women) and 7 heterozygous (2 men and 5 women).

For the *GSTP1* gene in exon 3, we found only 1 deletion in homozygous (1 woman). For exon 4 we do not found any alteration.

## 4 Discussion

Deletion in the *GST*s genes apparently results from unequal homologous exchanges involving flanking regions containing sequences with high identity that were identified as regions of deletion / addition of the null genotype [8, 9, 10, 11]. The homozygous deletion of the *GSTM1* gene is observed at frequencies ranging from 20 to 70% in different populations, while for gene *GSTT1* this variation is 11 to 38% [12, 13, 14].

The functional allele of *GSTM1* are active in detoxification of various substances, such as polycyclic aromatic hydrocarbons and other products of combustion, which are potentially genotoxic oxidizing agents [15], while *GSTT1* presents important activity in detoxification of peroxidized lipids and DNA oxidation products [16]. So that deletion of these isoforms has been associated with several types of diseases correlated with smoking, with emphasis on susceptibility studies to cancer [17, 18, 19, 20].

GSTP1 is a major enzyme metabolizing anticancer drugs like oxiplatin, cyclophosphamide which are used in the treatment of breast and colorectal cancer. An over expression of this enzyme causes resistance to drugs like cisplatin [21]. Therefore, investigation of these mutations will provide a clue to the investigation of responders to cancer therapy with certain chemotherapeutic drugs.

It appears that the absence of one or more forms of *GST* can become more susceptible to chemical and oxidative stress cell, which can lead to cell dysfunction, being of great importance to characterize the influence of deletion polymorphism *GSTM1* and *GSTT1* risk disease [3, 4].

In addition, pharmacogenetic has recently taken an important role showing significant differences, inter and intra population in metabolism, efficacy and toxicity of drugs, and this fact varies between regions studied. The Brazilian population, especially in the central-west region of Brazil offers a great research potential because it presents a unique admixture and diversity of variant combinations in different loci enabling gene interaction studies [22].

Deletion in the *GST*s genes apparently results from unequal homologous exchanges involving flanking regions containing sequences with high identity that were identified as regions of deletion / addition of the null genotype [8, 9, 10, 11]. The homozygous deletion of the *GSTM1* gene is observed at frequencies ranging from 20 to 70% in different populations, while for gene *GSTT1* this variation is 11 to 38% [12, 13, 14].

## 5 Conclusion

This study provides de first results of alterations distribution of GSTs M1, T1 and P1 mutations in a health population in Goiânia-Go, using the MLPa technic. Hence, it opens up new avenues for further investigations by epidemiologists in determining inter individual variation in genetic susceptibility to various diseases caused due to gene-environment interaction and the application of MLPA-based genotyping to routine clinical analysis will enable patients to be assigned to more accurate genotypes at a reasonable cost in a large number of individuals at the majority of locations.

## References

[1] Bhattacharjee S., Zhao Y., Hill J. M., Culicchia F., Kruck T. P., Percy M. E., et al. Selective accumulation of aluminum in cerebral arteries in Alzheimer’s disease. J. Inorg. Biochem. 126, 35–37. 2013.

[2] Sharma, A.; Pandey, A.; Sharma, S.; Chatterjee, I., Mehrotra, R.; Sehgal, A.; Sharma. J.K. Genetic polymorphism of glutathione S-transferase P1 (GSTP1) in Delhi population and comparison with other global populations. Meta Gene, 134–142, 2014.

[3] Wang, G., Zhang, L., Li, Q. Genetic polymorphisms ofGSTT1, GSTM1, andNQO1 genes and diabetes mellitus risk in Chinese population. Biochem. Biophys. Res. Commun. 341, 310–313. 2006.

[4] Zhang J, Liu H, Yan H, Huang G, Wang B. Null genotypes of GSTM1 and GSTT1 contribute to increased risk of diabetes mellitus: a meta analysis. Gene 518: 405–11. 2013.

[5] Iafrate AJ, Feuk L, Rivera MN, Listewnik ML, Donahoe PK, Qi Y, Scherer SW, Lee C. Detection of large-scale variation in the human genome. Nat Genet 36:949–951. 2004.

[6] Sebat J, Lakshmi B, Troge J, Alexander J, Young J, Lundin P, Maner S, Massa H, Walker M, Chi M, Navin N, Lucito R, Healy J, Hicks J, Ye K, Reiner A, Gilliam TC, Trask B, Patterson N, Zetterberg A, Wigler M, Large-scale copy number polymorphism in the human genome. Science 305:525–528. 2004.

[7] Cantsilieris, S., White, S.J., 2013. Correlating multiallelic copy number polymorphisms with disease susceptibility. Hum. Mutat. 34, 1–13.

[8] Xu, Q.; Lin, G.; Chen, W.J.; Zhou, X.; Lin, L. Relation between the expression of P-gp and GST-pi in oral and maxillofacial squamous carcinoma and chemoresistance, 37 (2002), pp. 90–93

[9] Sprenger, R.; Schlagenhaufer, R.; Kerb, R.; Bruhn, C.; Brockmöller, J.; Roots, I.; Brinkmann, U. Characterization of the glutathione S-transferase GSTT1 deletion: discrimination of all genotypes by polymerase chain reaction indicates a trimodular genotype-phenotype correlation. Pharmacogenetics, v. 10, n. 6, p. 557–65, 2000.

[10] Parl, F. F. Glutathione S-transferase genotypes and cancer risk. Cancer Letters, v. 221, p. 123–129, 2005.

[11] Reis, A.A.S.; Silva, D.M.; Curado, M.P. Da Cruz, A.D., Involvement of CYP1A1, GST, 72TP53 polymorphisms in the pathogenesis of thyroid nodules. Genet. Mol. Res. 9 (4): 2222–2229. 2010.

[12] ARRUDA et al., 1998

[13] BUFALO et al., 2006

[14] Bid HK, Konwar R, Saxena M, Chaudhari P, Agrawal CG, Banerjee M. Association of glutathione S-transferase (GSTM1, T1 and P1) gene polymorphisms with type 2 diabetes mellitus in north Indian population. J Postgrad Med 56: 176–181. 2010.

[15] Hayes, J.D., Pulford, D.J., 1995. The glutathione S-transferase supergene family: regulation of GST* and the contribution of the enzyme to cancer chemoprotection and drug resistance. Crit. Rev. Biochem. Mol. Biol. 30, 445–600

[16] Schneider, J., Bernges, U., Philipp, M., Woitowitz, H.J., 2004. GSTM1, GSTT1, and GSTP1 polymorphism and lung cancer risk in relation to tobacco smoking. Cancer Lett. 208, 65–74.

[17] Bell DA, Taylor JA, Paulson DF, Robertson CN, Mohler JL, Lucier GW. Genetic risk and carcinogen exposure: a common inherited defect of the carcinogen-metabolism gene glutathione S-transferase MI (GSTMl) that increases susceptibility to bladder cancer. J Natl Cancer Inst 1993: 85: 1159–1164.

[18] London, S. J.; Yuan, J. M.; Chung, F. L.; Gao, Y. T.; Coetzee, G. A.; Ross, R. K.; Yu, M. C. Isothiocyanates, glutathione S-transferase M1 and T1 polymorphisms, and lung-cancer risk: a prospective study of men in Shanghai, China. Lancet, v. 356, p. 724–729, 2000.

[19] Habdous, M.; Siest, G.; Herbeth, B.; Vincent-Viry, M.; Visvikis, S. Glutathione S-transferases genetic polymorphisms and human diseases: overview of epidemiological studies. Ann. Biol. Clin., v. 62, n. 1, p. 15–24, 2004.

[20] Lima-Jr., M. M.; Oliveira, M. N. L.; Granja, F.; Trindade, A. C. G.; De CASTRO SANTOS, L. E. M.; Ward, L. S. Lack of association of GSTT1, GSTM1, GSTO1, GSTP1 and CYP1A1 polymorphisms for susceptibility and outcome in brazilian prostate cancer patients. Folia Biologica(Praha), v. 54, p. 102–108, 2008.

[21] Salah, G.B. et al, Polymorphisms of glutathione S-transferases M1, T1, P1 and A1 genes in the Tunisian population: An intra and interethnic comparative approach. Gene 498 (2012) 317–322.

[22] C.T.X. Silva1,2, N.B. Costa1,2, K.S.F. Silva1,2, R.E. Silva1,3 and K.K.V.O. Moura1,2 Association between primary open angle glaucoma and genetic polymorphisms GSTM1/GSTT1 in patients from Goiônia Central-West Region of Brazil. Genetics and Molecular Research 13 (4): 8870–8875 (2014)

